# Constitutively active *SARM1* variants found in ALS patients induce neuropathy

**DOI:** 10.1101/2021.04.16.439886

**Authors:** A. Joseph Bloom, Xianrong Mao, Amy Strickland, Yo Sasaki, Jeffrey Milbrandt, Aaron DiAntonio

**Affiliations:** Needleman Center for Neurometabolism and Axonal Therapeutics and Department of Genetics, Washington University School of Medicine in Saint Louis, St. Louis, MO, USA; Needleman Center for Neurometabolism and Axonal Therapeutics and Department of Developmental Biology, Washington University School of Medicine in Saint Louis, St. Louis, MO, USA

**Keywords:** ALS, SARM1, neurodegeneration, axon, NAD

## Abstract

In response to injury, neurons activate a program of organized axon self-destruction initiated by the NAD^+^ hydrolase SARM1. In healthy neurons SARM1 is autoinhibited, but single amino acid changes can abolish autoinhibition leading to constitutively-active SARM1 enzymes that promote degeneration when expressed in cultured neurons. To investigate whether naturally-occurring human variants might similarly disrupt SARM1 autoinhibition and potentially contribute to risk for neurodegenerative disease, we assayed the enzymatic activity of 29 rare *SARM1* alleles identified among 8,507 amyotrophic lateral sclerosis (ALS) patients. Ten missense variants or small in-frame deletions exhibit constitutive NADase activity, including more than half of those that are unique to the ALS patients or that occur in multiple patients. Expression of these constitutively active ALS-associated SARM1 alleles in cultured dorsal root ganglion (DRG) neurons is pro-degenerative and cytotoxic. Intrathecal injection of an AAV expressing the common *SARM1* reference allele is innocuous to mice, but a construct harboring *SARM1*^*V184G*^, the constitutively active variant found most frequently in the ALS patients, causes axon loss, motor dysfunction, and sustained neuroinflammation. These results implicate rare hypermorphic *SARM1* alleles as candidate genetic risk factors for ALS and other neurodegenerative conditions.

## Main Text

Trauma and disease in the nervous system activate an intrinsic axon self-destruction pathway, also known as Wallerian degeneration, which facilitates the orderly clearance of damaged axon segments. This choice between maintaining or actively dismantling axons is primarily determined by the action of SARM1, a TIR-containing NAD^+^ hydrolase that cleaves NAD^+^ to generate nicotinamide and cyclic ADPR (cADPR), a useful biomarker of SARM1 activity^1^. In healthy neurons, SARM1 is maintained in an autoinhibited state, but injury- or disease-induced depletion of the axon survival factor NMNAT2 activates SARM1 leading to a rapid loss of NAD^+^, metabolic catastrophe, and axon fragmentation^2–4^. SARM1 knockout mice are viable and without apparent phenotypes under routine conditions, but are protected against neurodegeneration in models of axotomy, traumatic brain injury, peripheral neuropathy, glaucoma, and retinal degenerative diseases^5–11^. Conversely, mutations that decrease NMNAT2 activity lead to polyneuropathy in both humans and model organisms^12,13^, suggesting that aberrant SARM1 activation has a role in human disease. Furthermore, the recent observation that single point mutations in *SARM1* can disrupt enzyme autoinhibition^14–18^ led us to speculate that naturally-occurring human variants might similarly dysregulate SARM1 and thereby increase disease risk.

To investigate whether SARM1 mutations that disrupt its autoinhibition are associated with neurogenerative disorders, we sought to identify rare prodegenerative *SARM1* missense variants in human databases. ALS warrants particular attention because peripheral axon degeneration accompanies and may precede motoneuron death during ALS progression^19,20^. Here, we identify over two dozen such polymorphisms found in ALS patients and interrogate the activities of the encoded enzymes in cultured neurons. Provocatively, the majority of our strongest candidate variants disrupt SARM1 regulation and confer constitutive activity *in vitro*. Furthermore, expression of a constitutively-active *SARM1* allele found in three unrelated patients causes an ALS-like phenotype—motor dysfunction, axon loss and sustained neuroinflammation—when expressed in the mouse spinal cord via intrathecal delivery.

We identified a total of 29 *SARM1* coding variants (missense and small in-frame deletions) culled from three large publicly-accessible ALS consortia databases that include 8,507 cases in total (Table 1)^21–23^.

**Table 1.** Rare *SARM1* missense variants and in-frame deletions found in ALS patients.

| Variant | rsID | Percent Minor Allele Frequency<sup>a</sup> |  |  | Number of occurrences in ALS patients |
| --- | --- | --- | --- | --- | --- |
|  |  | African | East Asian | European<sup>b</sup> |  |
| Constitutively active |  |  |  |  |  |
| Δ226-232 | rs782325355 | 0 | 0 | 0.01 | 2<sup>d</sup> |
| Δ249-252 |  | 0 | 0 | 0 | 1<sup>d</sup> |
| V184G | rs71373646 | 0.007 | 0.006 | 0 | 3<sup>e</sup> |
| G206S | rs1555585199 | 0 | 0 | 0 | 2<sup>e</sup> |
| L223P |  | 0 | 0 | 0 | 1<sup>d</sup> |
| R267W | rs11652384 | 0 | 0 | 0.001 | 1<sup>e</sup> |
| V331E | rs1555585331 | 0 | 0 | 0 | 1<sup>d</sup> |
| E340K | rs781854217 | 0 | 0 | 0.003 | 1<sup>d</sup> |
| T385A |  | 0 | 0 | 0 | 1<sup>d</sup> |
| T502P | rs782421919 | 0 | 0 | 0.006 | 2<sup>d,f</sup> |
| Not constitutively active |  |  |  |  |  |
| A240E | rs1449836804 | 0 | 0 | 0.004 | 1<sup>d</sup> |
| R244S |  | 0 | 0 | 0 | 1<sup>d</sup> |
| A250T | rs1555585243 | 0 | 0.06 | 0 | 1<sup>d</sup> |
| A275V | rs376587698 | 0 | 0 | 0.006 | 1<sup>d</sup> |
| R310H | rs369186722 | 0 | 0 | 0.01 | 1<sup>d</sup> |
| A341V | rs373458416 | 0 | 0 | 0.01 | 2<sup>d</sup> |
| R403P | rs782706244 | 0 | 0 | 0.0009 | 1<sup>f</sup> |
| E431G | rs1555585662 | 0 | 0 | 0.001 | 1<sup>d</sup> |
| R465T |  | 0 | 0 | 0 | 1<sup>d</sup> |
| R484C | rs1555585809 | 0 | 0 | 0.0009 | 1<sup>d</sup> |
| A488E | rs782228906 | 0.004 | 0 | 0.02 | 2<sup>d</sup> |
| V518L | rs782106973 | 0 | 0 | 0<sup>c</sup> | 3<sup>d</sup> |
| R569C | rs571724138 | 0 | 0 | 0.005 | 1<sup>d</sup> |
| R570Q | rs539229444 | 0 | 0.008 | 0.005 | 1<sup>e</sup> |
| D637Y | rs1451417529 | 0 | 0 | 0 | 1<sup>d</sup> |
| A646S | rs782676389 | 0 | 0 | 0.0008 | 1<sup>d</sup> |
| M672V | rs782774927 | 0 | 0 | 0.004 | 2<sup>e,f</sup> |
| S684F | rs782256561 | 0.004 | 0 | 0.004 | 1<sup>f</sup> |
| R702C | rs781850558 | 0 | 0.005 | 0.002 | 1<sup>f</sup> |

<sup>a</sup>gnomAD v2
<sup>b</sup>non-Finnish
<sup>c</sup>V518L = 0.009% in gnomAD v3 non-Finnish Europeans
<sup>d</sup>MinE<sup>32</sup>
<sup>e</sup>ALS Variant Server<sup>34</sup>
<sup>f</sup>ALS Knowledge Portal<sup>33</sup>

Altogether, rare *SARM1* variants (i.e. with allele frequencies <0.01% in all gnomAD populations^24^) occur in >0.9% of ALS cases but in only ∼0.25% of controls^21,22^, and in only ∼0.3% of the general population^24^. For comparison, potentially pathogenic *TBK1* variants were reported in 1.1% of ALS cases and 0.19% of controls^25^. This suggests that rare pathogenic *SARM1* alleles might contribute to ALS risk.

To investigate whether these variants disrupt autoinhibition, we assayed the NAD^+^ hydrolase activities of the encoded enzymes. We prioritized the variants and first tested a) those identified in multiple ALS patients but not in healthy controls and b) those unique to ALS patients (i.e. not reported in any prior human study as of January 2020). These 15 *SARM1* variants (Figure 1A, Table 1) account for 51% (20/39) of rare *SARM1* variants in ALS patients genotyped in the three large ALS databases we investigated. To examine the properties of these mutants, we tested them in *Sarm1*^*−/−*^ mouse dorsal root ganglion DRG neurons. We prepared lentiviruses for the 15 *SARM1* mutant constructs, infected *Sarm1*^*−/−*^ neurons, and assessed their NAD^+^ hydrolase activity. Eight of these variants were determined to be constitutively active, i.e. we found that the baseline level of NAD^+^ was decreased in neurons expressing these mutant constructs and the level of cADPR, a specific *in vitro* and *in vivo* SARM1 biomarker^26^ was increased (Fig. 1). By contrast, *SARM1*^P332Q^, the only variant common in any gnomAD population (1.1% in Europeans) is not constitutively active (Fig 1).

**Figure 1.**
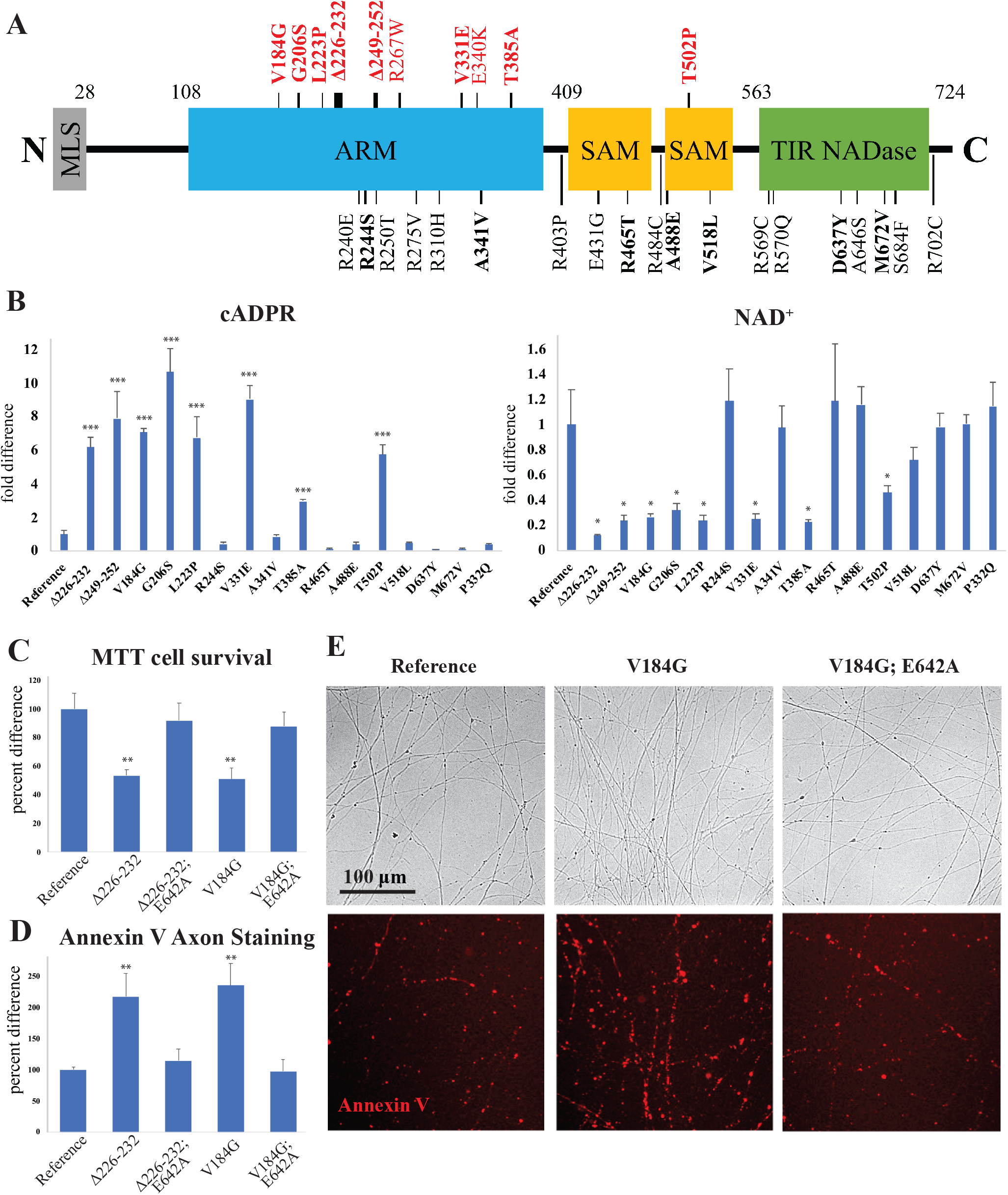
Dysregulated *SARM1* variants found in ALS patients promote neurodegeneration. **(A)** Schematic representation of the domain structure of SARM1. Constitutively-active variants are indicated above in red. Bold variants were prioritized because they were identified in multiple ALS patients or were unique to ALS patients. Δ indicates an in-frame deletion. MLS, mitochondrial localization signal; ARM, HEAT/Armadillo motif; SAM, sterile alpha motif; TIR, Toll/interleukin-1 receptor homology domain. **(B)** cADPR and NAD^+^ levels from cultured Sarm1^−/−^ DRG neurons infected with variant human *SARM1* constructs, performed in triplicate, relative to the reference *SARM1* allele. **(C)** Neuron death as measured by the MTT assay and **(D)** axon degeneration as measured by Annexin V staining in Sarm1^−/−^ DRG neurons infected with lentivirus expressing *SARM1* variant constructs as well as double mutant constructs including E642A, a point mutation that disrupts SARM1 NAD^+^ hydrolase activity, relative to the common *SARM1* reference allele, all performed in triplicate. **(E)** Representative bright-field and Annexin V-stained images of axons from Sarm1^−/−^ DRG cultures infected with variant and *SARM1* reference allele constructs. *p<0.05; **p<0.005; ***p<0.0005 difference from reference allele, two-tailed t-test.

Encouraged by these results, we assayed the activities of an additional 14 rare missense variants. These were considered poorer candidates because each is observed in only a single ALS patient and they are not unique to the patients as they are also found in the gnomAD database. Among these, we identified two additional constitutively active variants (Table 1). In total, 40% (4/10) of the SARM1 variants with constitutive NAD^+^ hydrolase activity occur in multiple ALS patients.

Point mutants that disrupt SARM1’s autoinhibitory interfaces result in dysregulation of SARM1 activity and promote the degeneration of cultured neurons^14–18^. Consistent with those findings, lentiviral-mediated expression of all the constitutively active variants we tested (Table 1) alters the cell body morphology of cultured *Sarm1*^*−/−*^ mouse DRG neurons consistent with cell death. To quantify this pro-degenerative activity, two variant constructs, *SARM1*^*V184G*^ and *SARM1*^*Δ226-232*^, were studied further. The mutant enzymes were expressed in *Sarm1*^*−/−*^ DRGs neurons and degeneration was measured by two methods. Fluorescently-labeled Annexin V, which binds to phosphatidylserine, was used to determine whether the expression of either variant construct significantly compromises axon health. Annexin V binding is a useful proxy for axon health as neurites undergoing Wallerian degeneration expose phosphatidylserine on their extracellular surfaces similarly to apoptotic cells^27^. Neuronal death was quantified using an oxidoreductase activity assay, a common measure of cell viability. Both assays demonstrated that both ALS-associated *SARM1* variants produced a significant degenerative effect relative to the common reference allele (Fig. 1).

While these variants exhibit constitutive NAD^+^ hydrolase activity, it is formally possible that they mediate their pro-degenerative effects via a distinct toxic mechanism. To investigate this alternate hypothesis, we generated constructs containing two mutations, either of the ALS-associated variants, *SARM1*^*V184G*^ or *SARM1*^Δ*226-232*^, together with E642A, a point mutation in the TIR domain that disrupts the catalytic glutamate required for SARM1 NAD^+^ hydrolase activity and axon degeneration^3^. In both cases, introducing E642A abolishes enzymatic activity and the detrimental effects of the constructs on cell body and axon health (Fig. 1). Hence, these ALS patient-derived *SARM1* variants promote degeneration via loss of autoinhibition and resulting constitutive NAD^+^ hydrolase activity.

To test whether the rare *SARM1* variants promote neurodegeneration *in vivo*, AAV viral vectors were administered intrathecally to male and female six-week old wild-type mice, expressing either the common human allele of *SARM1* (the reference allele) or *SARM1*^*V184G*^, the constitutively active variant found most frequently in the ALS patient databases. In these constructs, each SARM1 protein was fused to EGFP and expressed under the control of the human synapsin promoter. The AAV viruses were produced with these constructs and a mixture of PHP.S and PHP.eB serotype capsids (both derived from AAV9^28^) in order to infect neurons in the spinal cord and DRGs.

Animals injected with AAV expressing the common *SARM1* allele had no discernible behavioral phenotypes for at least 12 weeks. By contrast, those injected with AAV-*SARM1*^*V184G*^ exhibited motor impairment. Two of the mice rapidly progressed to full limb paralysis 3-4 days after injection. Other animals injected with *SARM1*^*V184G*^ (7/9) displayed less dramatic motor deficits characterized by hindlimb clasping^29^, and significant muscle weakness, as measured by the inverted screen assay (Fig. 2). These deficits were detected within 3 weeks of injection and did not progress significantly through the 12-week observation period.

**Figure 2.**
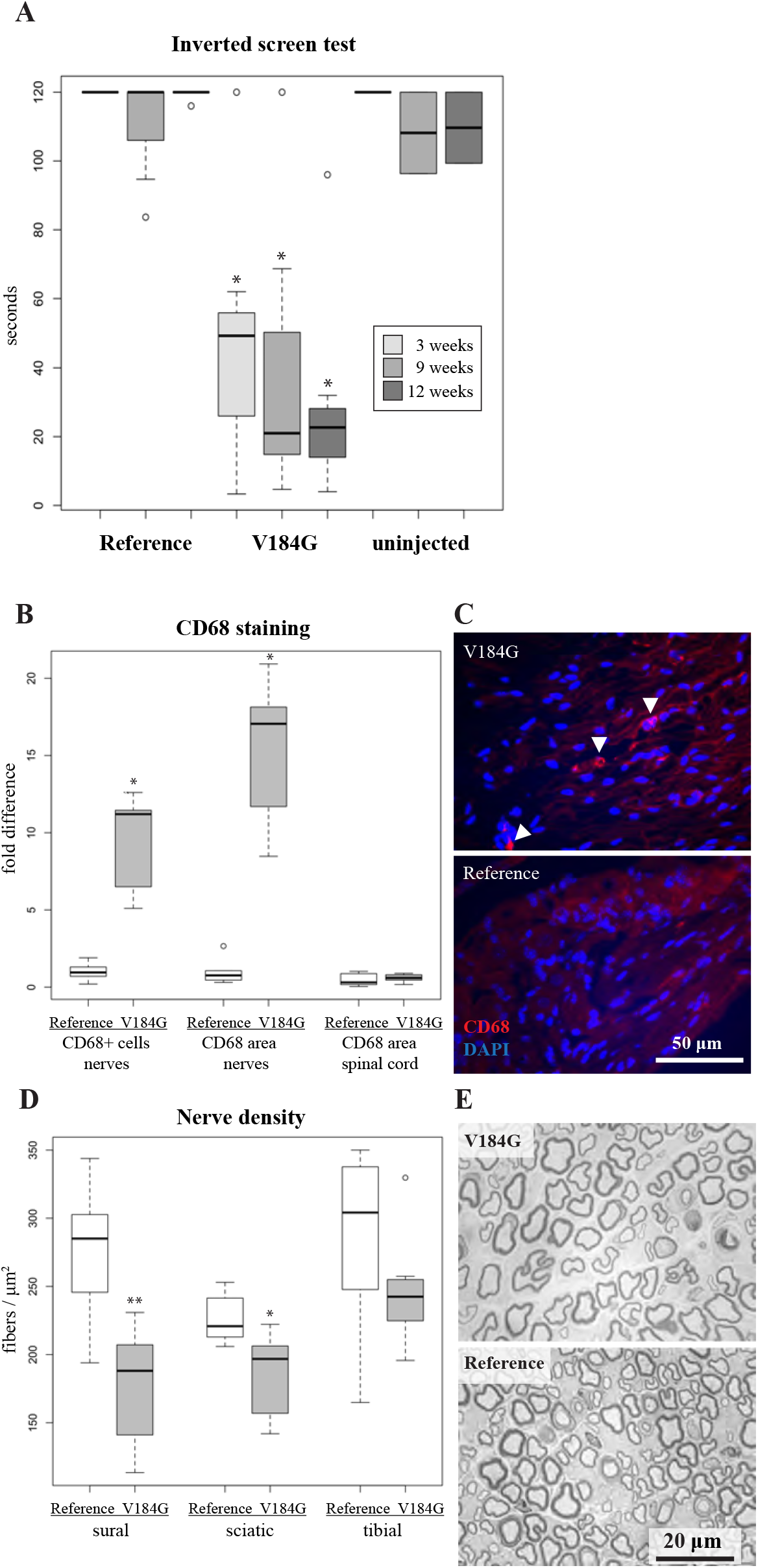
Motor dysfunction, neuroinflammation and axon loss in mice injected intrathecally with a *SARM1*^*V184G*^ AAV construct. **(A)** Average time suspended from an inverted screen (maximum 120 seconds, performed in triplicate) for C57/BL6 mice injected with a human *SARM1* reference allele (n=8) or *SARM1*^*V184G*^ (n=7) AAV compared to uninjected controls (n=3) 3, 9 and 12 weeks post-injection. *p<0.005 difference from both the reference allele and uninjected controls, 2-tailed t-test. **(B)** The normalized average number of cells stained by the macrophage marker anti-CD68 in nerve, and the average percent area of total anti-CD68 staining in nerve and in spinal cord sections, from C57/BL6 mice injected with a *SARM1*^*V184G*^ AAV construct (3 images per mouse, n=7 mice) relative to those injected with a human *SARM1* reference allele construct (n=8 mice) 12 weeks post-injection; *p<10^−4^, 2-tailed t-test. **(C)** Representative images of nerve stained with DAPI and anti-CD68 from mice 12 weeks after injection with a *SARM1*^*V184G*^ or reference allele construct. **(D)** Average fibers per cross-sectional μm^2^ in sural, sciatic and tibial nerves from mice 12 weeks after injection with a *SARM1*^*V184G*^ (3 images per mouse, n=7 mice) or reference allele construct (n=8 mice); *p<0.05; **p<0.001, 2-tailed t-test. **(E)** Representative images of toluidine blue stained sural nerve sections.

To characterize the neurodegeneration caused by *SARM*^*V184G*^ expression, the intrathecally injected mice were examined for evidence of axon degeneration and neuron loss. We examined the two mice that became rapidly paralyzed and the other mice with less severe disease as separate cohorts because of the difference in phenotype. In the spinal cords of the paralyzed mice, there was clear evidence of cell death around the ependymal canal as detected by TUNEL staining. Neuroinflammation was observed throughout the spinal cord of these mice as evidenced by prevalent staining for CD68, a marker of activated macrophages. Neither of these phenotypes were observed in animals injected with the common *SARM1* allele construct (Additional File 1). These mice did not display obvious myelin defects or vacuolization in the sural, sciatic or tibial nerves at 3-4 days post-infection.

The mice treated with *SARM1*^*V184G*^ that displayed a less severe behavioral response were sacrificed twelve weeks post-injection. Pathological inspection of their spinal cords revealed no evidence of ongoing apoptosis or elevated CD68 staining. Their peripheral nerves, however, contain almost 10-fold more CD68-positive macrophages than those treated with the control *SARM1* allele (Fig. 2). Macrophages also increase in size upon activation^30^, and the *SARM1*^*V184G*^-infected mice have a 1.6-fold greater CD68-stained area per cell than do control mice, yielding a 15.2-fold difference in total CD68-stained area. Hence, neuronal expression of *SARM1*^*V184G*^ triggers an elevated inflammatory response in peripheral nerves that persists for at least twelve weeks after treatment^31^.

The average fiber densities in the peripheral nerves of the *SARM1*^*V184G*^-injected mice are also lower than those injected with AAV expressing the common human allele, demonstrating that expression of the constitutively active SARM1 promotes axon loss. The number of axons were counted in the sural, sciatic and tibial nerves. In both the sural and sciatic nerves, the density of axons is significantly lower (p<0.001), a trend that is evident in the tibial nerve, though it did not reach statistical significance (Fig. 2). The average axon size and extent of myelination (g-ratio) does not differ significantly between the variant and common *SARM1* allele treated animals (p>0.1, n=15). Myelin defects and vacuolization are not observed in these nerves, indicating a lack of ongoing axon loss. The lack of axon defects at twelve weeks is consistent with the early, but stable, deficits in motor function observed in mice receiving the *SARM1*^*V184G*^ virus (Fig. 2). We interpret these data as evidence that a subset of neurons—those sufficiently susceptible to SARM1-dependent degeneration and infected with virus—lost their axons before three weeks, while others, including uninfected neurons, remained healthy and functional up to twelve weeks. Inter-animal differences in motor dysfunction severity likely reflect variability in infection efficiency.

In summary, we find that many rare *SARM1* variants found in ALS patients also lack normal autoinhibition, and that such an allele induces neurodegeneration and neuroinflammation when expressed in the mouse nervous system. We therefore propose that hypermorphic *SARM1* mutations are a candidate congenital risk factor for ALS. The mechanism by which constitutive NAD^+^ hydrolase activity would predispose to neurodegeneration appears straightforward as low NAD^+^ is a death sentence for energy-hungry neurons and is associated with both disease and aging-related functional defects^32^. We speculate that the contrast between virus-transfected mice that rapidly display severe degenerative phenotypes, and human ALS patients who are typically diagnosed only after several decades of life, likely reflects the difference in SARM1 expression—i.e. viral over-expression precipitates abrupt metabolic catastrophe in this model, whereas chronic suboptimal NAD^+^ levels lead to gradual motoneuron attrition in patients. Fortunately, small molecule SARM1 inhibitors are already in development^33^, and we have shown that a *SARM1* dominant negative gene therapy can potently block the SARM1 programmed axon degeneration pathway in mice^34^. Establishing that SARM1 inhibition is safe and effective in carriers of pathogenic *SARM1* variants could provide a vital stepping stone toward the use of SARM1-directed therapeutics more generally for ALS and other diseases that involve axon degeneration.

## Methods

### Mice

Male and female WT and *Sarm1* knockout C57/BL6 mice were housed and used under the direction of institutional animal study guidelines at Washington University in St. Louis. The inverted screen test of strength was performed as previously^35^. All protocols received institutional IACUC approval.

### DRG culture

Mouse DRG culture was performed as previously described^36^. DRG were dissected from embryonic day 13.5 *Sarm1* knockout C57/BL6 mouse embryos and cells suspended in growth medium at a concentration of ∼7 × 10^6^ cells/ml in 96-well tissue culture plates (Corning) coated with poly-d-Lysine (0.1 mg/ml; Sigma) and laminin (3 μg/ml; Invitrogen). For axotomy, suspended neurons (2 μl) were placed as a drop in the center of each well and incubated at 37°C with 5% CO_2_ for 15 min, after which media was added to each well. Lentiviral particles containing *SARM1* variants were generated as previously described^36^. Lentivirus was added (1– 10 × 10^3^ pfu) after 1–2 days (DIV) and metabolites were extracted or assays were performed at 6–7 DIV. Cell death was quantified by assaying mitochondrial function (MTT assay), as previously described^37^.

### Automated quantification of axon degeneration

The axon degeneration index, the ratio of fragmented axon area over total axon area, was quantified as previously described^36^. To quantify Annexin V staining, the Alexa Fluor(tm) 568 conjugate (ThermoFisher) was added to the cultured neurons at a 1:100 dilution four days after viral infection. Bright field and fluorescent images were acquired one hour later using Operetta. Unbiassed image analysis was performed using ImageJ as follows: total axon area was measured from the binary bright field images after subtracting background. For Annexin fluorescent intensity measurement, the fluorescent images were background subtracted and then annexin positive area was defined using the particle analyzer. Data was reported as the total fluorescent intensity of the annexin positive area divided by the axon area.

### DRG metabolite extraction and metabolite measurement

At DIV6, tissue culture plates were placed on ice and culture medium replaced with ice-cold saline (0.9% NaCl in water, 500 μl per well). Saline was removed and replaced with 160 μl ice cold 50% MeOH in water. Solution was transferred to tubes containing 50 μl chloroform on ice, shaken vigorously, and centrifuged at 20,000*g* for 15 min at 4 °C. The clear aqueous phase (140 μl) was transferred into microfuge tubes and lyophilized under vacuum. Lyophilized samples were reconstituted with 5 mM ammonium formate (15 µl), centrifuged (13,000 *g*, 10 min, 4°C), and 10 µl of clear supernatant was analyzed. NAD^+^ and cADPR were measured using LC-MS/MS as previously described^26^.

### AAV constructs and virus injections

AAV particles with a mixture of Php.s and Php.eB capsids^28^, containing a human *SARM1* gene construct fused to EGFP, under the control of the human synapsin promoter, were produced by the Viral Vector Core of the Hope Center for Neurological Disorders at Washington University in St. Louis. Viral particles were purified by iodixanol gradient ultracentrifugation and virus titers were measured by dot blot. Under light anesthesia with Avertin, 6 × 10^11^ viral genomes were injected intrathecally at L6/S1. Viral expression in mice 12-weeks post injection was confirmed by detecting EGFP expression via immunohistochemical analysis of sectioned DRGs.

### Immunohistochemistry, imaging and quantification

After perfusion with PBS followed by 4% PFA in PBS, tissues were fixed in 4% PFA in PBS for 1 h at room temperature and placed in 30% sucrose in PBS overnight at 4°C, then embedded in OCT (Tissue-Tek), frozen on dry ice, and then stored at −80°C. Longitudinal sections of 6 μm or cross-sections of 20 μm were obtained using a cryostat and slides were stored at −20°C. DRG and nerve slides were post-fixed in cold acetone, then washed with PBS. Spinal cord slides were simply washed three times in PBS. All slides were subsequently blocked with 4% BSA and 1% Triton X-100 in PBS and incubated with rat anti-CD68 (1:500; Bio-Rad) and mouse-anti-GFP conjugated to Alexa Fluor 488 (1:250; Thermo Fisher Scientific) overnight in the blocking buffer. Slides were then washed, incubated in secondary antibodies (Jackson ImmunoResearch Laboratories) and mouse anti-GFP conjugated to AlexaFlour 488 (1:250, Thermo Fisher Scientific), washed, and mounted in Vectashield with DAPI. Slides were imaged using a DMI 4000B confocal microscope (Leica Microsystems) with a 20× oil objective and DFC 7000-T camera (Leica Microsystems). For quantification, at least four images were measured per animal. CD68-positive cells were counted by a researcher blinded to the images’ treatment group. The total CD68-stained area and nerve area in each image was quantified with the particle analyzer in ImageJ using a uniform threshold.

### TUNEL apoptosis detection

TUNEL was performed as previously described^38^. Slides prepared for immunohistochemistry were thawed then postfixed with 4% PFA for 10 min at room temperature, washed thoroughly with PBS, incubated with 10 μg/ml proteinase K for 15 min at 37°C, then washed with PBS. A positive control slide was incubated in DNase I (1 U/ml) for 1 h at RT, then washed with PBS. Slides were then pretreated with TdT buffer (25 mm Tris-HCl, 200 mm sodium cacodylate, 0.25 mg/ml BSA, 1 mm cobalt chloride, Roche Diagnostics) at RT for 10 min. To perform end-labeling, TdT buffer was combined with terminal deoxynucleotidyl transferase (Roche Diagnostics, 400 U/slide) and Biotin-16-dUTP (Roche Diagnostics, 4 μm) and added to slides for 1 h at 37°C. Slides were thoroughly washed with PBS, then blocked for 30 min with 5% normal goat serum in PBS with 0.3% Triton-X, then incubated with Alexa-Fluor-conjugated streptavidin (Jackson ImmunoResearch Laboratories) for 30 min at 37°C. Slides were washed and mounted in Vectashield with DAPI.

### Toluidine blue staining and axon quantification

Sural, sciatic and tibial nerves were fixed in 3% glutaraldehyde in 0.1 M PBS, processed and imaged as previously described^8^. Micrographs were stitched using Leica software and axons were counted using ImageJ. To determine axon size distribution and G ratios of the sciatic nerve, four nonoverlapping areas per cross section were imaged with a 100× oil objective of a Zeiss Axioskop and photographed with a Hitachi camera. Photographs were analyzed using a previously described binary imaging analysis method^39^.

### Statistical analysis

Two-tailed significance is reported throughout. All statistics were calculated using the R software package. All data is available upon request.

## Abbreviations

AAV: Adeno-associated virus
ALS: Amyotrophic Lateral Sclerosis
ARM: HEAT/Armadillo motif
cADPR: Cyclic adenosine diphosphate ribose
CD68: Cluster of Differentiation 68
DRG: Dorsal root ganglion
EGFP: Enhanced green fluorescent protein
MLS: Mitochondrial localization signal
MTT: 3-(4,5-dimethylthiazol-2-yl)-2,5-diphenyltetrazolium bromide
NAD: Nicotinamide adenine dinucleotide
NMNAT2: Nicotinamide mononucleotide adenylyltransferase 2
TIR: Toll/Interleukin receptor
TUNEL: Terminal deoxynucleotidyl transferase dUTP nick end labeling
SAM: Sterile alpha motif
SARM1: Sterile alpha and TIR motif containing 1

## Declarations

### Ethical Approval and Consent to participate

All studies were approved by the Washington University Institutional Animal Care and Use Committee.

### Consent for publication

Not Applicable

### Availability of supporting data

All data relevant to this study are contained within the article.

### Authors’ contributions

A.J.B, J.M and A.D designed the study. X.M performed in vitro experiments. A.S performed in vivo experiments. Y.S assisted with mass spec analysis and method development. A.J.B. analyzed the data. A.J.B., J.M. and A.D. drafted and edited the figures and manuscript. All authors read and approved the final manuscript.

## Acknowledgements

We extend our thanks to Tim Fahrner and Kelli Simburger for cloning viral constructs, to Rachel McClarney and Cassidy Menendez for assistance with mouse husbandry and administering behavioral tests, and to Alicia Neiner for processing LC-MS/MS samples. This work was supported by the Hope Center Viral Vectors Core at Washington University School of Medicine.

## Authors’ information

Not Applicable

## Figure Legends

**Additional file 1.**
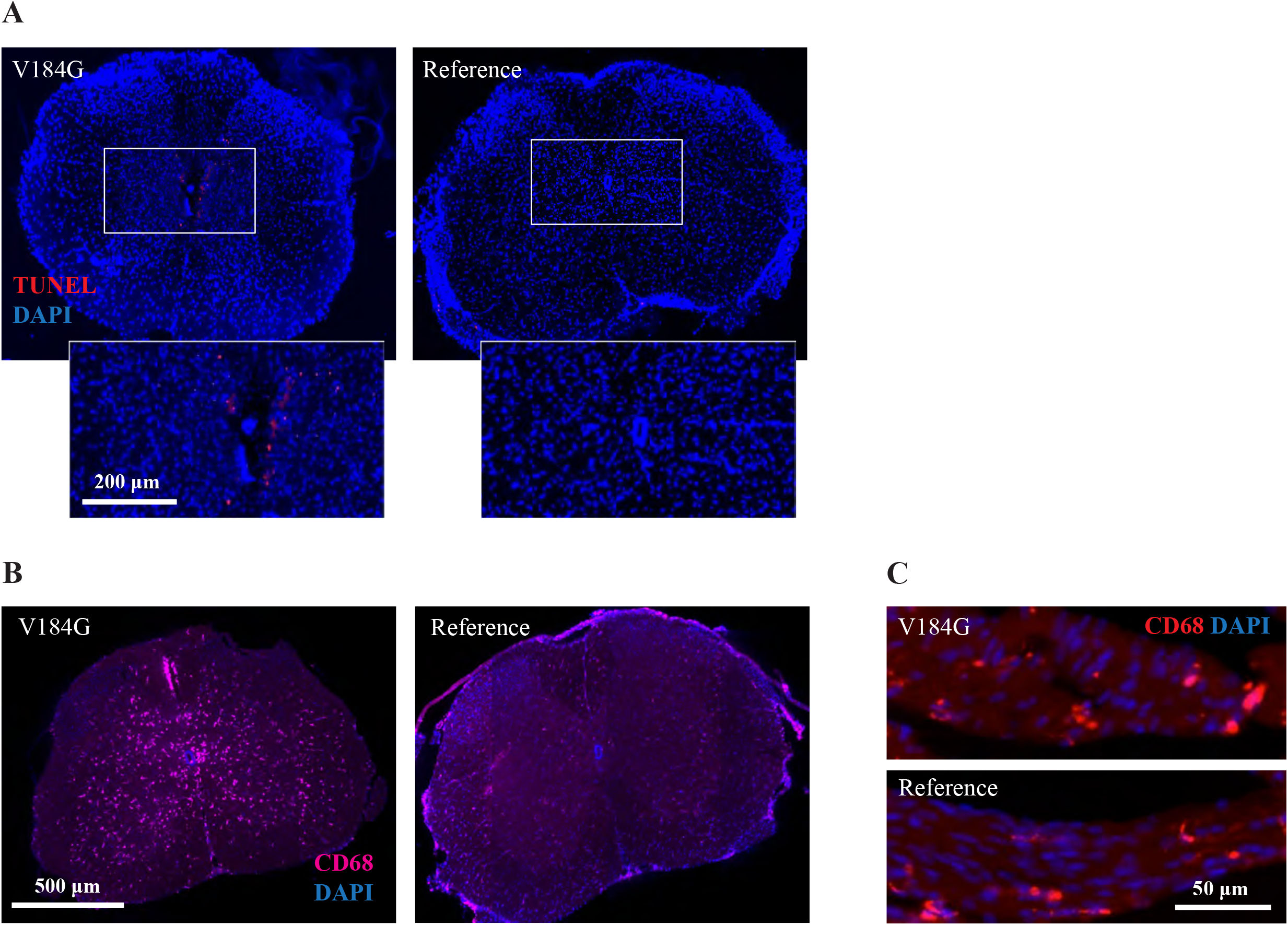
Rapid cell death and neuroinflammation in mice injected intrathecally with a *SARM1*^*V184G*^ AAV construct. **(A)** Representative images of spinal cord sections, with closeup of ependymal canal, stained with DAPI and the apoptosis marker TUNEL from mice 2 days after injection with a *SARM1*^*V184G*^ or *SARM1* human reference allele construct. **(B)** Representative images of spinal cord and **(C)** adjacent nerve sections stained with DAPI and the macrophage marker anti-CD68 from mice 2 days after injection with a *SARM1*^*V184G*^ or reference allele construct.

